# Transcriptional consequences of herpes simplex virus 1 ICP4 inducible expression in uninfected cells

**DOI:** 10.64898/2025.12.09.693233

**Authors:** Nora L Herzog, Sarah Keegan, Rebecca Plessel, Liam J Holt, Ian J Mohr, Angus C Wilson

## Abstract

Successful viral infections reflect the balanced outcome of a tightly regulated program of viral gene expression and manipulation of the host cell environment to favor production of new infectious particles. The productive (lytic) replication cycle of herpes simplex virus 1 (HSV-1) is dependent on the essential transcription factor ICP4 encoded by one of five immediate-early genes. At different steps in the HSV-1 temporal cascade, ICP4 either positively or negatively regulates transcription of immediate-early, early, and late HSV-1 genes, largely through sequence-specific binding to cis-acting regulatory elements. In contrast, the direct regulation of host transcription by ICP4 is less well understood. In this study we exogenously expressed doxycycline-inducible wild-type and mutant ICP4 proteins in uninfected primary human fibroblasts and performed RNA-Seq to identify ICP4-driven changes to the host transcriptome. Cross-referencing our findings to a published dataset of ICP4-dependent changes to the host transcriptome in HSV-1 infected cells provided validation for a subset of differentially-expressed genes regulated by ICP4. Furthermore, disrupting the ICP4 DNA-binding domain was sufficient to alter the cellular gene transcriptional program responsive to ICP4. This indicates that the DNA-binding domain of ICP4, which is required for site-specific DNA binding to the virus genome, may also regulate binding to the host genome. Together, these data provide a comprehensive transcriptomic analysis of how wild-type and mutant ICP4 expression impact cellular gene expression in uninfected cells.

**IMPORTANCE:** Herpes simplex virus 1 (HSV-1) is a widespread human pathogen responsible for a variety of mild to severe disease states. The viral lifecycle is driven by the essential viral transcription factor ICP4, which ensures the coordinated expression of most viral genes. Expression of ICP4-regulated genes and viral lifecycle progression impacts host gene expression; therefore, determining the specific impact of ICP4 on host gene expression has been challenging. Here, we provide a more direct analysis, examining changes in the transcriptome of primary fibroblasts when ICP4 is expressed alone. Differentially expressed genes were identified and cross-referenced to published data from infections with wild-type and ICP4-null viruses. We show that mutations in ICP4 that disrupt interactions with the host transcriptional machinery or interfere with site-specific DNA binding further modified this ICP4-driven remodeling of the host transcription. These data provide a comprehensive transcriptomic analysis of ICP4-driven dysregulation of host gene expression in uninfected cells.

## INTRODUCTION

As obligate intracellular parasites, viruses have evolved to manipulate cellular functions and the tissue microenvironment to drive their own life cycles. Herpes simplex virus type 1 (HSV-1) is an example of an extremely successful viral pathogen that has evolved alongside humans for millennia, infecting at least 60% of the world’s population(1, 2). HSV-1 is a large linear double-stranded DNA virus in the Herpesviridae family(3). These viruses alternate between productive (lytic) and latent infections. Productive (lytic) replication is characterized by high expression of viral proteins, amplification of the viral genome, and production of new infectious progeny(3). Following primary infection of the epithelium, HSV-1 establishes latency in neurons of the peripheral nervous system leading to a lifelong infection, the most common consequences of which are oral and genital blisters associated with episodic reactivation of latent virus(4).

The HSV-1 genome encodes for at least 84 viral proteins and several noncoding RNAs(3). These genes are divided into four kinetic classes, immediate-early (IE, 𝛼), early (E, 𝛽), leaky-late (𝛾_1_), and true-late (𝛾_2_)(3, 5). Transcription of the E genes, which primarily involve functions necessary for amplification of the HSV-1 genome, requires the IE protein ICP4 (𝛼4/IE175/Vmw175), one of five proteins produced during the initial stage of each infection cycle(3, 5). ICP4 is a large 175 KDa (1298 residue) phosphoprotein that can oligomerize on DNA and is found predominantly in the nucleus of infected cells(3, 6, 7). It is an essential transcriptional activator for both E and late gene transcription, while also acting as a transcriptional repressor of IE genes(3, 7–13). In addition to binding to viral promoter sequences, ICP4 also binds promiscuously to host DNA through early phases of HSV-1 infection, though this binding is later reduced(13). ICP4 has also been shown to interact with cellular transcription machinery, including RNA Pol II and its associated factors, as well as chromatin remodelers, including the ATPase subunits of SWI/SNF, INO80, and NuRD complexes, to stimulate viral gene transcription(7, 9, 13–18). Additionally, ICP4 is thought to be responsible for the preferential recruitment of RNA Polymerase II (RNAPII) to the HSV-1 genome throughout infection, leading to a substantial reduction in RNAPII occupancy across the host genome(13, 19, 20).

Functioning as a robust and versatile transcription factor, ICP4 both activates and represses transcription of HSV-1 genes through site-specific DNA binding to cognate target sites in the virus genome as well as through more generalized binding(20, 21). Three major functional domains have been described: the N-terminal activation domain (NTA), the C-terminal activation domain (CTA), and the DNA-binding domain (DBD)(22). Sequence-specific binding conferred by the DBD is required but is not sufficient to induce viral gene expression during infection(15, 23). Transcriptional activation is achieved by recruiting general transcription factor complexes including TFIID and Mediator, as well as RNAPII(15, 17, 18, 24, 25). These interactions are mediated by the intrinsically disordered regions in the N-terminal activation domain and the DNA-binding domain of the ICP4 protein(25, 26). Taken together, this suggests that ICP4 may function to regulate host cell transcription as well as viral transcription during HSV-1 infection.

While the role of ICP4 in regulating viral transcription has been well studied, how ICP4 impacts host transcription during HSV-1 infection has only been evaluated by comparing cells infected with wild-type ICP4 or ICP4 mutant HSV-1 viruses. Bulk RNA-Seq on human MRC5 fibroblasts infected with either wild-type (WT) or ICP4-null viruses together with genome-wide ChIP-Seq have identified host genes that are differentially regulated during HSV-1 infection in an ICP4-dependent manner(13, 20, 27). However, the extent to which these changes in gene expression reflect the specific action of ICP4 alone or instead require other ICP4-dependent activities in uninfected cells has not been previously addressed.

To investigate how ICP4 alone influences host gene expression in uninfected cells, we constructed lentiviral vectors to express WT or mutant ICP4 variants from a doxycycline-inducible promoter, independent of HSV-1 infection. The ICP4 mutants evaluated have been previously described in the literature and correspond to truncations lacking the first 90 amino acids of the N-terminus (ΔNTA), lacking the entire C-terminal activation domain (ΔCTA), and a variant containing a double amino acid substitution in the DNA-binding domain (mDBD) that disrupts site-specific binding to the canonical ICP4 responsive sequence upstream of the ICP4 gene but is not thought to prevent more general binding to host chromatin(7, 22, 28–31). While global transcription rates were not detectably altered, bulk RNA-sequencing showed that a distinctive transcriptional program resulted from inducible ICP4 expression in uninfected primary human fibroblasts. Mutations in N- and C-terminal activation domains and the site-specific DNA binding domain interfered with these ICP4-induced transcriptional changes. Our data reveals how inducible ICP4 expression in uninfected cells impacts host gene transcriptional programming. Moreover, it suggests that the site-specific DNA-binding domain of ICP4, which mediates virus genome recognition, may also regulate binding to the host genome and contribute to reprogramming host gene expression.

## RESULTS

### Controlled expression of ICP4 in uninfected primary fibroblasts

To directly address if and how ICP4 impacts cellular gene expression in the absence of an ongoing productive HSV-1 infection, we constructed doxycycline (Dox)-inducible lentiviruses to express either wild-type ICP4 (WT) or mutant ICP4 variants. Three previously characterized ICP4 mutants were chosen for this analysis: a deletion of the C-terminal transactivation domain or CTA (ΔCTA, sequence adapted from n208 virus)(32), a truncation of the first 90 amino acids of the N-terminal transactivation domain or NTA (ΔNTA, sequence adapted from n6 virus)(32), and a double amino acid substitution mutation in the DNA-binding domain or DBD (mDBD, containing substitutions R456L-R457L)(28)(**Fig. 1A**). Following lentivirus transduction into primary human dermal fibroblasts (NHDFs), cells were treated + / - Dox for 9 h, after which total protein lysates were collected and the steady state ICP4 protein levels were determined by immunoblotting (**Fig. 1B**). Appropriate nuclear localization of WT and mutant ICP4 proteins was verified by indirect immunofluorescence (**Fig. 1C**). Taken together, these data established that all four inducibly-expressed ICP4 proteins accumulated to similar levels and were predominantly localized within the nucleus.

**Figure 1:**
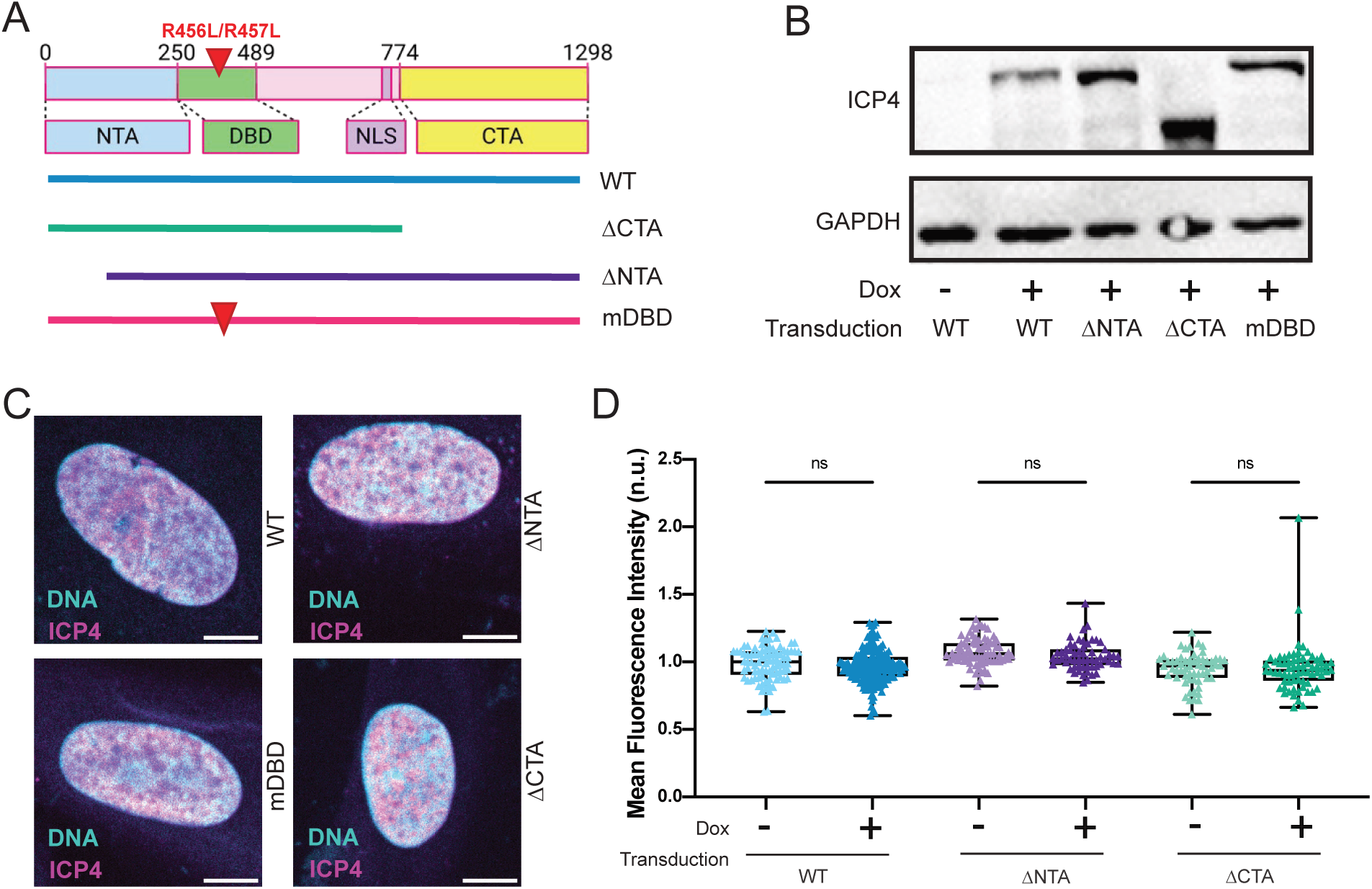
Global transcription rates in uninfected fibroblasts are not detectably altered by inducible ICP4 expression. (A) Schematic showing the major functional domains of WT ICP4 (WT) and boundaries of ICP4 mutants used in this study. The site-specific DNA-binding domain (DBD), nuclear localization signal (NLS), N-terminal transactivation domain (NTA) and C-terminal transactivation domain (CTA) are indicated. In addition to ICP4 N- and C-terminal truncation mutants (ΔNTA and ΔCTA, respectively), a double arginine to leucine substitutions at residues 456 and 457 within the DBD reported to disrupt sequence-specific DNA binding (mDBD) was also evaluated. (B) Immunoblot showing ICP4 protein accumulation after lentivirus transduction of NHDFs and Dox induction (9 h) for the different ICP4 constructs shown in (A). Blots are representative of N=3 biological replicates. (C) Representative indirect immunofluorescence (IF) images detecting ICP4 (*magenta*) in nuclei and DNA (SiR-DNA, *cyan*) from lentivirus-transduced, Dox-induced (9 h) NHDFs (scale bar, 10 μm). Data is representative of N=3 biological replicates. (D) NHDFs were transduced and treated -/+ Dox as in (B, C). Cells were exposed to the nucleoside analog 5-ethynyl uridine (5EU) for 2 h beginning at 7 h post treatment (hpt). At 9 hpt, transduced cells were fixed, processed for click chemistry, and probed by IF to detect ICP4. 5EU incorporation in ICP4-positive cells was scored as mean fluorescence intensity (MFI) using Foci-Counting and normalized to background. n>99; N≥3 biological replicates. All statistical comparisons were performed using a Kruskal-Wallis test. * signifies p<0.05, **** signifies p<0.0001.

### ICP4 expression does not alter global transcription levels

To determine whether ICP4 induction impacted global transcription rates, NHDFs transduced with each lentivirus underwent puromycin selection prior to dox-induction of ICP4 and 5-ethynyl uridine (5EU) exposure. Incorporation of 5EU into newly synthesized RNA was subsequently visualized in fixed cells using click chemistry to conjugate the alkyne group to an azide-modified fluorescent dye(33, 34). The intensity of fluorescence is proportional to the total amount of RNA produced during the 5EU pulse. Incorporation of 5EU was not detectably altered by induction of WT or mutant ICP4 proteins compared to the uninduced controls (**Fig. 1D**). Thus, expression of WT or mutant ICP4 is insufficient to detectably alter global rates of RNA biogenesis in uninfected NHDFs.

### Expression of ICP4 in uninfected cells invokes a distinctive transcriptional program

While global levels of RNA biogenesis were not detectably altered by inducible ICP4 expression, it remains possible that ICP4 alters the transcription of specific cellular RNAs. To investigate whether ICP4 expression can reprogram host cell gene expression in uninfected cells, Illumina short-read RNA-sequencing (RNA-Seq) was performed on polyA-selected RNA isolated from NHDFs that were either mock-transduced or transduced with lentiviruses that express WT or mutant ICP4 in response to doxycycline addition. An average of 84 million reads were obtained for each replicate. A principal components analysis (PCA) plot of the ICP4 induction condition compared to the mock transduced primary NHDFs showed a clear separation of the mock samples from the ICP4 induced samples (**Fig. 2A**). Normalized ICP4 expression levels as determined by the number of reads mapping to the ICP4 cassette for each sample used were analyzed and found some small level of variation across samples was found, which could potentially contribute to the dispersed appearance of the ICP4 induced samples on the PCA plot (**Fig. 2B**). However, when normalized ICP4 expression levels across all mutants were examined, comparable levels of ICP4 mRNA were observed across experiments with different mutants (**Fig. 2C**). To further validate these results, we generated a PCA plot following batch-correction for WT ICP4, ΔCTA, ΔNTA, and mDBD induction experiments, and found that each condition was largely clustered together (**Fig. 2D**). Batch effect could not be modeled for our mock condition due to a lack of matching induced genotypes from the sequencing run. However, examining PCA for each mutant with mock samples showed 86-89% variance on PC1, the axis that separated mock from the mutant genotypes. WT ICP4 samples were sequenced with each mutant ICP4 condition and showed 70% variance accounted for on PC1 (**Fig. 2D**), which separated mock samples from WT ICP4 samples. This was most likely due to expression differences of WT ICP4 and/or technical differences from performing separate sequencing runs. While it is possible that these technical differences are detected in differential expression (DE) analysis, given that the largest variance was detected on the PC axis separating mock from ICP4-induced genotypes, we considered the differences observed to reflect true biological signal.

**Figure 2:**
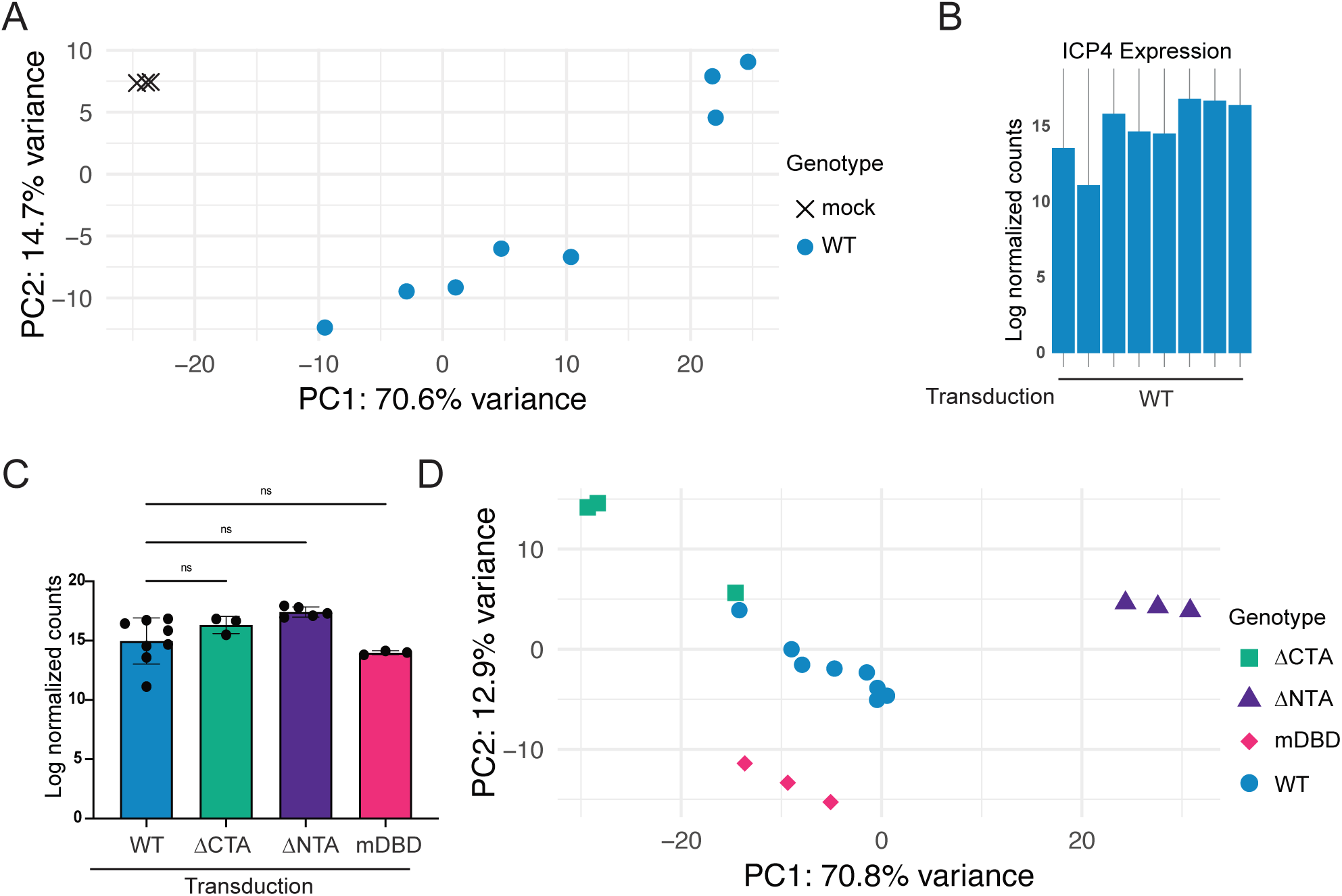
Induced ICP4 expression selectively remodels the transcriptome of uninfected fibroblasts. (A) PCA plot following batch-correction of WT ICP4-induced and mock RNA-Seq samples. RNA was collected from lentivirus-transduced NHDFs following 9 h of Dox induction. (B) Log-normalized counts for ICP4 in each replicate of WT induction for RNA-Seq data. (C) Pooled log normalized counts for ICP4 across replicates in each induction condition (WT, ΔCTA, ΔNTA, mDBD). (D) PCA plot following batch-correction of WT induced, ΔNTA induced, ΔCTA induced, and mDBD induced RNA-Seq samples.

Characterization of the NHDF transcriptome with or without WT ICP4 induction. This analysis revealed a set of differentially expressed genes (DEGs) when compared to the mock condition and is visualized by volcano plot using a log2FC threshold of 2 or -2 in addition to an adjusted p-value (padj) significance level of 0.01 (**Fig. 3A**). To further classify DEGs by gene ontology (GO) in terms of molecular function (GO MF, **Fig. 3B**) or biological process (GO BP, **Fig. 3C**), gene set enrichment analysis (GSEA) for pathway enrichment was performed(35). In our analysis, rank was determined by multiplying p-value by the sign of the log2FC, which ranks the most significant upregulated genes at the top of the list and the most significant downregulated genes at the bottom of the list. Following WT ICP4 induction, 606 genes were significantly upregulated (log2FC ≥2, padj ≤0.01) and 273 genes were significantly downregulated (log2FC ≤-2, padj ≤0.01). Downregulated genes were enriched in the GO terms *Catalytic activity acting on DNA, ATP dependent activity acting on DNA,* and *Helicase activity* as well as *DNA replication, Double strand break repair,* and *DNA recombination* (**Fig. 3B**, **Fig. 3C**). Leading edge genes, meaning the subset of genes that contribute most to the enrichment signal, within these GO terms included components of the chromatin remodeling complexes Ino80, SWI/SNF, and NuRD, as well as components of TFIID (**Table S1**), which have been found to co-purify with ICP4(9).

**Figure 3:**
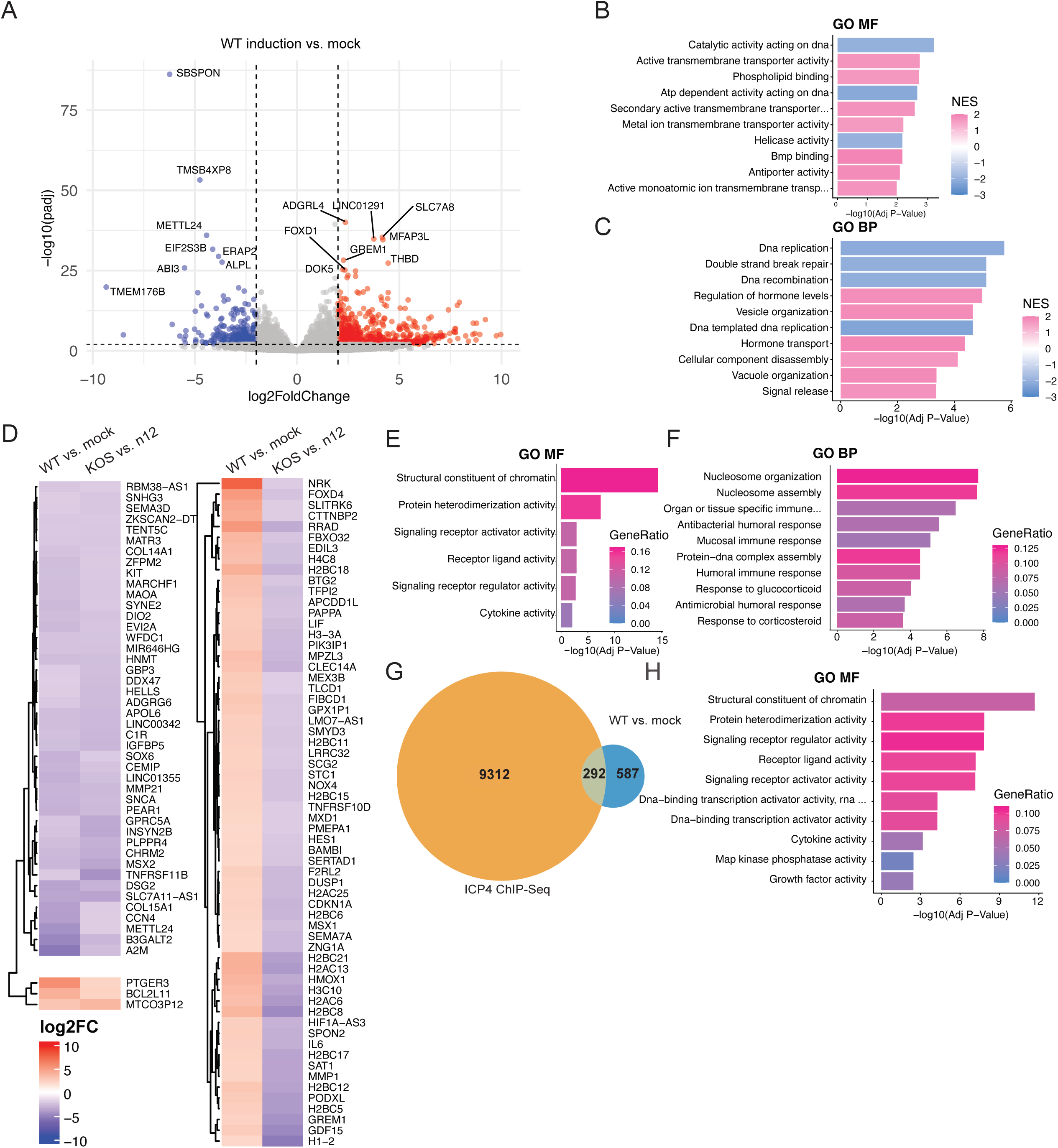
ICP4 induction reprograms the host transcriptome. (A) Volcano plot showing all differentially expressed genes (DEGs) in WT ICP4 induction compared to mock. X-axis represents log2FC with a threshold of 2 for upregulated genes (*red, leftmost dashed line*) or -2 for downregulated genes (*blue, rightmost dashed line*). Y-axis represents the -log10 of the adjusted p-value, with a threshold ≤0.01 (horizontal dashed line). The 8 upregulated and downregulated genes with the most significant adjusted p-values are labeled. (B, C) Gene set enrichment analysis (GSEA) for pathway enrichment on all genes with adjusted p-value ≤0.05 and log2FC ≥ |2|. Rank was determined by multiplying p-value by the sign of the log2FC, which ranks the most significant upregulated genes at the top of the list and the most significant downregulated genes at the bottom of the list. GSEA identifies pathways whose genes are enriched near the top or bottom of the list, calculating statistics by permutations of the gene sets. Top 10 most enriched pathways for molecular function GSEA (B) and biological process GSEA (C) are displayed. Normalized enrichment score (NES) heatmap indicates the degree to which the identified pathway was upregulated (pink) or downregulated (blue) in the given gene set. (D) Heatmap showing log2FC for DEGs identified using thresholds as in (A) (left column), compared to the log2FC of host genes calculated by comparing WT HSV-1 infection (KOS) to ICP4-null mutant HSV-1 infection (n12) at 12 hpi in MRC5 fibroblasts. KOS vs. n12 data was taken from publicly available RNA-Seq dataset PRJNA851702(56). Only genes with log2FC ≥ |2| in the KOS vs. n12 dataset are pictured. (E) Molecular function and (F) biological process GSEA as in (B, C) of genes identified as differentially expressed in both datasets: WT induction vs. mock = log2FC ≥ |2|, padj≤0.01; KOS vs. n12 = log2FC ≥ |2|. Gene ratio heatmap indicates the ratio of input genes that are present in the indicated pathway over the total number of input genes. (G) Venn diagram showing the overlap between all DEGs identified using thresholds as in (A) (WT vs. mock) and ICP4 host binding sites within the promoter & 5’UTR regions during HSV-1 infection at 2 hpi (ICP4 ChIP-Seq). ICP4 binding sites were identified using publicly available ICP4 ChIP-Seq dataset PRJNA553563(13). (H) Top 10 most enriched pathways from molecular function overrepresentation analysis (ORA) of identified overlapping genes in (G).

We then compared this RNA-Seq data to a published RNA-Seq analysis (<u>PRJNA851702</u>) of human MRC5 fibroblasts infected for 12 h with either WT HSV-1 (strain KOS) or the ICP4-null n12 virus (ΔICP4)(36, 37). A heat map comparing the log2FC of genes identified as differentially expressed during WT ICP4 induction (|log2FC| ≥ 2, padj≤0.01) to the log2FC of genes identified as differentially expressed in an ICP4-dependent manner (KOS vs. n12, |log2FC| ≥ 2) was generated (**Fig. 3D**). Three populations of genes were identified: those that are upregulated in both WT induction and KOS infection (3 genes, **Fig. 3D bottom left**); those that are downregulated in both WT ICP4 induction and KOS infection (53 genes, **Fig. 3D top left**); and those that are upregulated in WT ICP4 induction but downregulated in KOS infection (63 genes, **Fig. 3D right**) (**Table S2**). These three populations were assessed together in a single analysis using GSEA for pathway enrichment (**Fig. 3E**, **Fig. 3F**). Significant enrichment for the molecular function GO terms *Structural constituent of chromatin* and *Protein heterodimerization activity* (**Fig. 3E**) as well as biological process GO terms *Nucleosome organization, Nucleosome assembly* in addition to pathways involved in the immune response was observed (**Fig. 3F**).

Finally, 292 overlapping genes were identified by comparing DEGs in response to WT ICP4 induction in uninfected NHDFs with a published ICP4 ChIP-Seq dataset obtained from HSV-1 infected MRC5 fibroblasts at 2 hpi (<u>PRJNA553563</u>)(13) (**Table S3, Fig. 3G**). Only a small percentage of the genes identified by ICP4 ChIP-Seq from HSV-1 infected fibroblasts were found to be differentially expressed by WT ICP4 induction in uninfected fibroblasts (292 of 9604 identified binding sites). GO enrichment analysis for the overlapping genes(38) showed cellular component enrichment for the nucleosome as well as molecular function enrichment for *Structural constituent of chromatin, Protein heterodimerization activity*, and several pathways involved in receptor signaling and transcription activation (**Fig. 3H**).

### ICP4 N- and C-terminal transactivation domains impact transcriptional regulation of the host genome

To evaluate whether DEGs resulting from WT ICP4 induction were sensitive to changes in the ability of ICP4 to interact with the general transcription machinery, we performed RNA-seq on cells expressing each of the previously defined ICP4 truncation mutants.

Induction of ΔNTA expression identified a set of DEGs relative to the mock condition. (**Fig. 4A**). 1265 genes were significantly upregulated (log2FC ≥2, padj ≤0.01) and 485 genes were significantly downregulated (log2FC ≤-2, padj ≤0.01) by this N-terminal truncation. GSEA revealed that downregulated genes were enriched in the GO terms *Catalytic activity acting on DNA, ATP dependent activity acting on DNA, Helicase activity*, and *Single stranded DNA binding* (**Fig. 4B**) as well as *DNA replication, Double strand break repair, DNA recombination,* and *Regulation of chromosome organization* (**Fig. 4C**). Comparing the sets of DEGs resulting from ΔNTA with WT ICP4 induction revealed a significant overlap. Unexpectedly, ΔNTA produced a much larger cohort of DEGs, although many of these genes were uniquely responsive to ΔNTA (**Fig. 4D**). Interestingly, however, similar GO terms for molecular function (**Fig. 4E**) and biological process (**Fig. 4F**) were identified with both WT ICP4 and the ΔNTA truncation. Pathways uniquely enriched in response to ΔNTA induction included those enriched in the overlap of WT ICP4 induction compared to HSV-1 infection (**Fig. 3E**) and ICP4 ChIP-Seq (**Fig. 3H**), with enrichment for molecular function GO terms such as *Structural constituent of chromatin, Protein heterodimerization activity*, and several pathways involved in receptor signaling and transcription activation (**Fig. 4E**). However, biological processes involved in development, maturation, and differentiation were significantly and uniquely enriched in our subset of DEGs in response to ΔNTA induction (**Fig. 4F**). Lastly, we compared DEGs from ΔNTA induction with the promoter and 5’UTR binding sites from the ICP4 ChIP-Seq dataset and identified 604 overlapping genes (**Table S3**, **Fig. 4G**).

**Figure 4:**
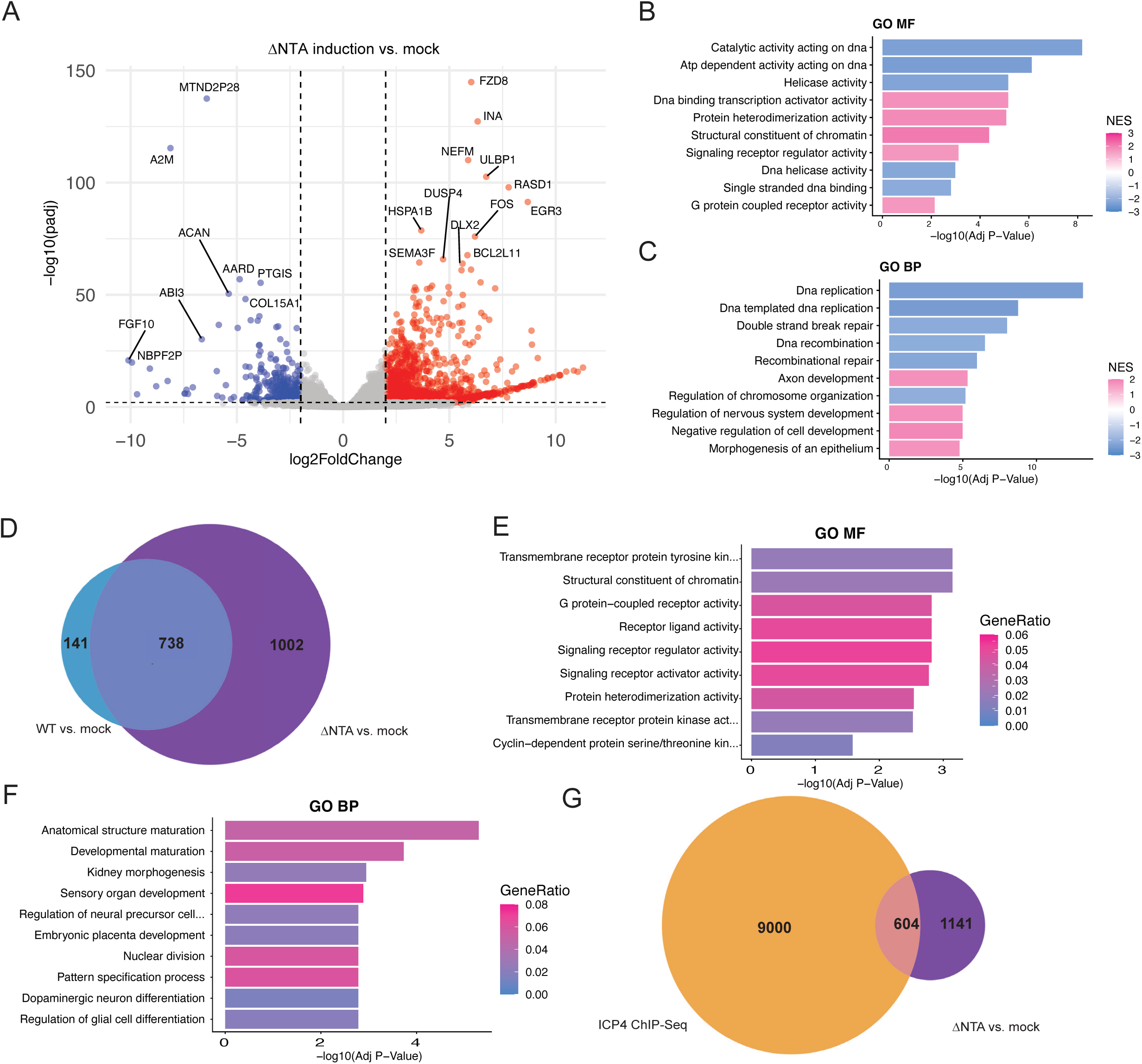
Mutations in the N-terminus interfering with ICP4 transcriptional activation alter transcriptomic responses in uninfected cells. (A) Volcano plot showing all differentially expressed genes (DEGs) in ΔNTA induction compared to mock, as in Fig. 2A. (B, C) Gene set enrichment analysis (GSEA) for pathway enrichment on all genes with adjusted p-value ≤0.05 and log2FC ≥ |2| as previously described. Top 10 most enriched pathways for molecular function GSEA (B) and biological process GSEA (C) are displayed. NES heatmap indicates the degree to which the identified pathway was upregulated (pink) or downregulated (blue) in the given gene set. (D) Venn diagram showing the overlap between all DEGs identified in **Fig 2A** (WT vs. mock) compared to the DEGs identified in (A) (ΔNTA vs. mock). (E) Top 10 most enriched pathways from molecular function ORA and (F) biological process ORA of DEGs identified as specific to ΔNTA (D, purple). Gene ratio heatmap indicates the ratio of input genes that are present in the indicated pathway over the total number of input genes. (G) Venn diagram showing the overlap between all DEGs identified in (A) (ΔNTA vs. mock) and ICP4 host binding sites within the promoter & 5’UTR regions during HSV-1 infection at 2 hpi (ICP4 ChIP-Seq). ICP4 binding sites were identified as in Fig. 2G.

To investigate differential gene expression during induction of the ΔCTA truncation, we compared our ΔCTA dataset to the mock condition and visualized these data by volcano plot as previously described (**Fig. 5A**). During ΔCTA induction, 442 genes were significantly upregulated (log2FC ≥2, padj ≤0.01) and 434 genes were significantly downregulated (log2FC ≤-2, padj ≤0.01). However, no enriched pathways were detected when we attempted to classify these genes in terms of molecular function. A reduced set of pathways were identified for genes that were upregulated, and these included the GO terms *Negative regulation of intracellular signal transduction*, *Signal release*, *Regulation of secretion*, *Hormone transport*, *Embryo implantation*, and *Vacuole organization* (**Fig. 5B**). Interestingly, enrichment pathways were not identified for downregulated genes. Comparing the set of DEGs identified during ΔCTA induction with those identified during WT ICP4 induction revealed a smaller cohort of overlapping genes than those identified during ΔNTA induction (**Fig. 5C**). The Venn diagram describing the overlap of DEGs confirmed the enriched genes in ΔCTA induction were distinct from those seen in WT ICP4 induction. While both *Hormone transport* and *Vacuole organization* were present in the enriched pathways during WT ICP4 induction, the overall transcriptional program during ΔCTA induction was significantly different. We attempted to classify genes differentially regulated only upon ΔCTA induction in terms of both molecular function and biological process, but pathway enrichment was not found for this subset of DEGs. As a final step, DEGs from ΔCTA induction were compared with the promoter and 5’UTR binding sites from the ICP4 ChIP-Seq dataset and 198 overlapping genes were identified (**Table S3**, **Fig. 5D**).

**Figure 5:**
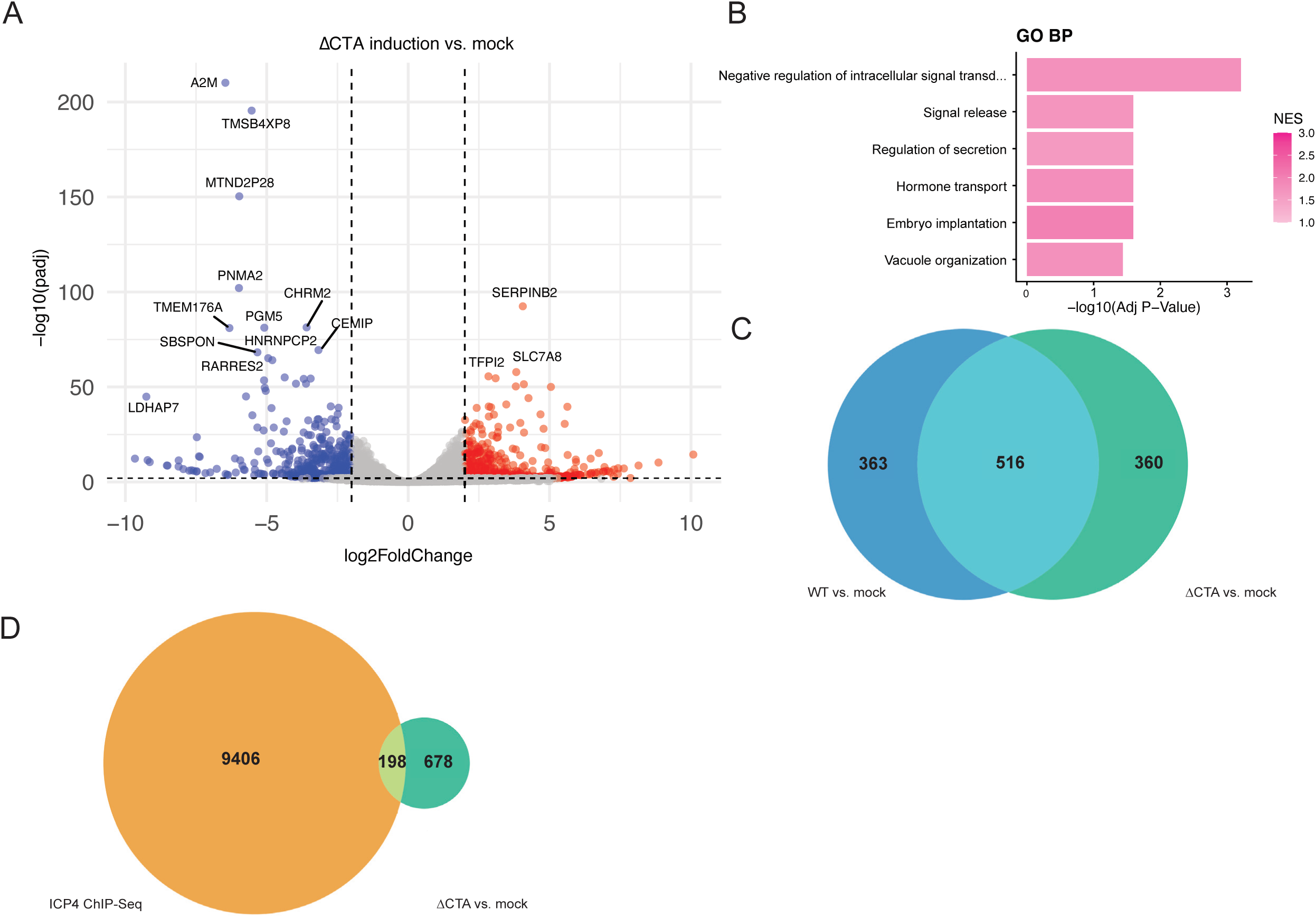
Mutations in the C-terminus interfering with ICP4 transcriptional activation alter transcriptomic responses in uninfected cells. (A) Volcano plot showing all differentially expressed genes (DEGs) in ΔCTA induction compared to mock as in Fig. 2A. (B) Gene set enrichment analysis (GSEA) for pathway enrichment on all genes with adjusted p-value ≤0.05 and log2FC ≥ |2| as previously described. The only 6 enriched pathways for biological process GSEA are displayed. No pathways for molecular function GSEA were identified. NES heatmap indicates the degree to which the identified pathway was upregulated (pink) or downregulated (blue) in the given gene set. (C) Venn diagram showing the overlap between all DEGs identified in **Fig 2A** (WT vs. mock) compared to the DEGs identified in (A) (ΔCTA vs. mock). No pathway enrichment was identified for DEGs specific to ΔCTA (C, teal). (D) Venn diagram showing the overlap between all DEGs identified in (A) (ΔCTA vs. mock) and ICP4 host binding sites within the promoter & 5’UTR regions during HSV-1 infection at 2 hpi (ICP4 ChIP-Seq). ICP4 binding sites were identified as in Fig. 2G.

### Impairment of ICP4 site-specific DNA binding modifies the host transcriptomic response

To investigate whether site-specific DNA binding activity was important for transcriptomic responses induced by ICP4 in uninfected cells, we expressed the mDBD ICP4 variant. The mDBD variant is deficient for specific binding to ICP4-regulated viral promoters *in vitro*, but has not been reported to alter non-specific binding of ICP4 to host chromatin(21, 22, 28, 29). Comparison of DEGs detected upon mDBD induction compared to the mock condition showed 401 significantly upregulated genes (log2FC ≥2, padj ≤0.01) and 587 significantly downregulated genes (log2FC ≤-2, padj ≤0.01) (**Fig. 6A**). The overall number of DEGs is considerably smaller compared to the amount observed following WT ICP4 induction, suggesting that the mDBD double amino acid substitution may be disrupting binding of ICP4 to cellular DNA. Downregulated genes were enriched in the GO terms *Helicase activity, Chromatin binding, Single stranded DNA binding*, and several other pathways involved in ATP activity and ATP-dependent activity on DNA (**Fig. 6B**) as well as biological process pathways involved in DNA replication, recombination, repair, and cell cycle maintenance (**Fig. 6C**). When comparing DEGs induced in response to mDBD with those found during WT ICP4 induction, significant differences were observed, underscoring the impact that the site-specific DNA-binding domain may have in regulating ICP4 binding to the host genome (**Fig. 6D**). Specifically, when assessing pathway enrichment among genes differentially expressed only upon mDBD induction, we found further enrichment of pathways involved in DNA replication and repair, cell cycle processes, and ATP-dependent activity (**Fig. 6E**, **Fig. 6F**).

**Figure 6:**
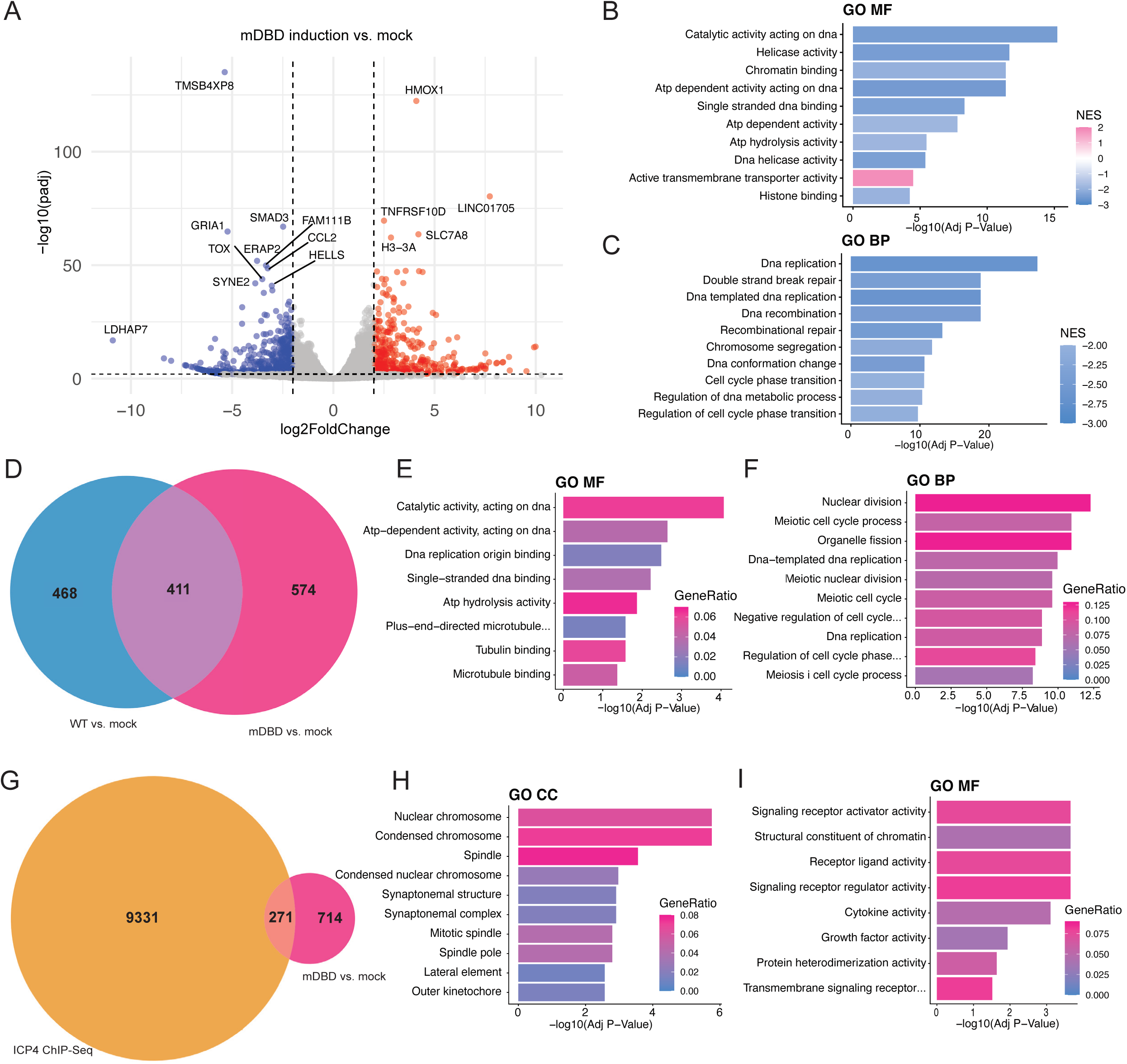
Mutation impairing ICP4 site-specific binding *in vitro* interferes with transcriptomic reprogramming in uninfected cells. (A) Volcano plot showing all differentially expressed genes (DEGs) in mDBD induction compared to mock as in Fig. 2A. (B, C) Gene set enrichment analysis (GSEA) for pathway enrichment on all genes with adjusted p-value ≤0.05 and log2FC ≥ |2| as previously described. Top 10 most enriched pathways for molecular function GSEA (B) and biological process GSEA (C) are displayed. NES heatmap indicates the degree to which the identified pathway was upregulated (pink) or downregulated (blue) in the given gene set. (D) Venn diagram showing the overlap between all DEGs identified in **Fig 2A** (WT vs. mock) compared to the DEGs identified in (A) (mDBD vs. mock). (E) Top 10 most enriched pathways from molecular function ORA and (F) biological process ORA of DEGs identified as specific to mDBD (D, hot pink). Gene ratio heatmap indicates the ratio of input genes that are present in the indicated pathway over the total number of input genes. (G) Venn diagram showing the overlap between all DEGs identified in (A) (mDBD vs. mock) and ICP4 host binding sites within the promoter & 5’UTR regions during HSV-1 infection at 2 hpi (ICP4 ChIP-Seq). ICP4 binding sites were identified as in Fig. 2G. (H) Top 10 most enriched pathways from cellular component and (I) molecular function overrepresentation analysis (ORA) of identified overlapping genes in (G).

Finally, comparison of DEGs from mDBD induction in uninfected NHDFs with a publicly available ICP4 ChIP-Seq dataset(13) from HSV-1 infected cells at 2hpi revealed 271 overlapping genes (**Table S3, Fig. 6G**). GO enrichment analysis on the overlapping genes showed cellular component enrichment for the nuclear chromosome and spindle (**Fig. 6H**) as well as molecular function enrichment for the many of the same pathways seen in WT ICP4 induction, including *Structural constituent of chromatin, Protein heterodimerization activity,* and several pathways involved in receptor signaling and transcription activation (**Fig. 6I**).

## DISCUSSION

ICP4 is an essential virus-encoded transcription factor that promotes the expression of the HSV-1 E and late genes, while downregulating HSV-1 IE genes, during the productive replication cycle. This is achieved in large part through site-specific DNA binding to cis-acting elements distributed across the HSV-1 genome(3, 21). The importance of ICP4 in regulating the HSV-1 life cycle is well established(8, 21, 36, 39). Whether ICP4 plays a significant role in regulating the expression of host genes is less clear because prior work has not directly examined the impact of ICP4 expression on the host transcriptome in the absence of other HSV-1 gene products. Instead, prior profiling experiments have either identified genes differentially regulated during HSV-1 infection in an ICP4-dependent manner or defined sites of ICP4 association in host chromatin during infection.

By cross-referencing these existing datasets with the DEGs we identified by profiling the transcriptomes of uninfected NHDFs with or without inducible ICP4 expression, changes in host gene expression brought about through the direct activities of ICP4 in the absence of other HSV-1 proteins can be revealed. One significant caveat is that while both the MRC5 cells used in the prior studies, and the NHDFs used in this study are human fibroblasts, they are from different individual donors. It is thus likely there are small differences in genome sequence and more importantly, differences in the epigenetic landscape of the chromosomal loci. Regardless, a detailed understanding of how ICP4 might directly alter the transcriptional landscape of the host cell provides fresh insight into its activity early in infection, before ICP4 is recruited to the newly replicated viral genomes.

The inclusion of the ICP4 mutants ΔNTA, ΔCTA, and the mDBD double missense mutant provides additional depth to this study. Firstly, transcriptomic studies using RNA-Seq on RNA from cells infected with either the n6 (ΔNTA) or n208 (ΔCTA) viruses have not been reported. Additionally, a recombinant mutant virus carrying the mDBD ICP4 mutation has not been constructed(28). By contrast, the capacity of ICP4 mutants lacking fully functional N-terminus and C-terminus domains to bind and recruit transcription machinery, including components of TFIIB, TFIID, and Mediator, has been examined in detail(7, 9, 15, 25, 40, 41). Specifically, various N-terminus truncation mutants including one analogous to n6 ICP4 showed reduced interaction with components of the general transcription machinery and a deficiency in negative regulation(7, 25, 41), while n208 ICP4 was found to be compromised for higher-level oligomerization of ICP4 homodimers on DNA(16, 29) as well as recruitment of specific subunits of ICP4-interacting transcription machinery(9, 15). Our analysis has now identified differences in the host transcriptional programs induced directly by these ICP4 mutants in uninfected fibroblasts.

First, induction with ΔNTA led to enrichment of genes in pathways distinct from those observed upon WT ICP4 induction (**Fig. 3B-C**, **Fig. 4B-C**) and greater upregulation of cellular transcripts than other ICP4 constructs (**Fig. 4A**). This latter observation suggests that the N-terminal 90 amino acids of ICP4 may be important for transcriptional repression, as ICP4 is known to act as a transcriptional repressor and activator(15, 20, 21, 24).

Second, ΔCTA induction showed significantly less overlap between DEGs seen during induction and ICP4-ChIP-Seq peaks compared to other ICP4 constructs, with a total of 198 overlapping genes (**Fig. 5D**). Given that the ΔCTA ICP4 isoform lacks robust transcription factor activity(7, 29), this in itself is not surprising. However, DEGs upon ΔCTA induction were resistant to classification, showing no enrichment of molecular function pathways and only a small subset of weakly upregulated pathways in the GO term “biological processes” (**Fig. 5B**). It is possible that in the absence of the C-terminal activation domain, ICP4 binds more weakly and randomly to host chromatin, creating an unclassifiable pattern of gene expression in contrast to when there is strong, specific binding to gene sets.

Third, though the mDBD ICP4 mutant protein was not previously thought to alter host DNA binding, clear differences in cellular gene expression resulting from WT ICP4 and mDBD induction were observed, with less than half of the respective identified DEGs being shared (**Fig. 2D**, **Fig. 6D**). In particular, upregulation of pathways involved in transporter activity in WT ICP4 induction compared to mDBD induction illustrates that even this small alteration in the site-specific DNA-binding domain leads to differential gene expression. It may be worthwhile to re-examine the importance of ICP4 binding motifs for the binding of ICP4 to host genomic sites, which has been thought to be largely nonspecific and determined primarily by accessibility(13, 22, 28, 29).

Finally, with the exception of ΔCTA, gene expression changes induced by all ICP4 variants in uninfected fibroblasts showed enrichment for pathways involving DNA replication, DNA recombination, nucleosome organization, and chromosome/DNA remodeling. While HSV-1 is known to marginalize and condense host chromatin late in infection, ICP4 has not been directly linked to this phenomenon(42, 43). However, ICP4 mediates the recruitment of transcription machinery to replicating viral genomes(13, 17, 18), and this recruitment has recently been implicated in the remodeling of host chromatin during HSV-1 infection(44). Furthermore, ICP4 has been shown to copurify with components of chromatin remodeling complexes, but the nature and functional consequences of these interactions remains elusive. Recently, inhibition of the SWI/SNF chromatin remodeling complex was shown to lead to a loss of chromatin accessibility and a decrease in transcriptional activity(45). Additionally, SWI/SNF activity has also been implicated in alleviating nucleosome formation during promoter-proximal pausing (PPP)(46), a process that is important for regulating HSV-1 transcription and thought to be controlled at least partly by ICP4 activity(20, 47, 48). During HSV-1 infection, the transcriptional environment is competitive between host and viral genes; ICP4 has been identified as a factor responsible for weighing the transcriptional environment in favor of the virus(13). Collectively, these findings raise the possibility that ICP4 may be acting on a transcriptional level to control the remodeling of the host chromatin and the overall nuclear environment.

While additional studies are needed, our findings suggest that ICP4 activity in the absence of other HSV-1 encoded components can produce nuanced transcriptional outputs in uninfected cells that might be influenced by DNA sequence, chromatin context, or other preexisting host factors. The availability of this new dataset exploring host transcriptional reprogramming responsive to inducible ICP4 expression in uninfected cells will help advance our understanding of how host transcriptional regulation may be altered or controlled during HSV-1 infection.

## ACKNOWLEDGEMENTS

AW was supported by the following NIH grants: R01-AI176335 and R01-AI170583. IM was supported by NIH grants: R01-GM056927, R01-AI073898, and R01-AI176335. LJH was supported by The Hypothesis Fund https://doi.org/10.13039/100031064 and a National Institutes of Health (NIH) award R01-GM132447.

We thank the NYU Langone Genome Technology Center (RRID: SCR 017929), in particular the help of Peter Meyn and Shyanon Rai in library preparation and sequencing. The Genome Technology Center is supported by the Cancer Center Support Grant P30CA016087. We would also like to thank all members of the Holt, Mohr, and Wilson labs for their feedback, advice, and encouragement.

## SUPPLEMENTAL MATERIAL

**Table S1** Gene ontology (GO) enrichment terms including leading edge genes identified during WT ICP4 induction compared to mock.

**Table S2** Populations of overlapping genes identified in WT ICP4 induction compared to mock and WT HSV-1 infection compared to mutant n12 infection.

**Table S3** Information on transcripts identified during WT and mutant ICP4 induction and their overlap with ICP4 ChIP-Seq peaks in promoter and 5’UTRs at 2 hpi.

## MATERIALS & METHODS

### Data Availability

All sequencing data are publicly accessible in the SRA database; sequencing dataset generated for this study can be found at PRJNA1368599; existing sequencing datasets can be found at PRJNA851702 and PRJNA553563. Visualizations of data and comparisons can be accessed through RShiny app, Herp-IE-dia: http://159.89.47.161/icp4_shiny/.

### Cell Culture

Normal human dermal fibroblasts (Lonza; CC-2509) were cultured in Dulbecco’s modification of Eagle’s medium (DMEM) (Corning; 10-013-CV) supplemented with 10% (v/v) fetal bovine serum (FBS) (Gemini bio-products; 100-106) and 100 U/mL penicillin-100 μg/mL streptomycin (Corning; MT-30-002-Cl). Cells were grown at 37°C and 5% CO_2_.

### Lentivirus stock preparation and transduction

HEK293T cells (9×10^6^ per 15-cm dish) were plated in antibiotic-free DMEM (Gibco, Cat. No. 11995073) supplemented with 10% FBS. The next day, cells were transfected with a mixture of the appropriate lentiviral vector plasmid together with packaging plasmids (psPAX2 and pMD2.G), using fuGENE HD transfection reagent (Promega, Cat. no. E2312) following manufacturer’s protocol. After 24 h the media was replaced with antibiotic-free DMEM and supernatants collected at 72 h post-transfection, aliquoted and stored at -80°C. Infectious lentivirus was introduced into NHDFs via reverse transduction with 5 mL of virus stock in fresh media and 10 µg/mL polybrene (Sigma-Aldrich; TR-1003-G), replacing the media after 24 h. Cells were selected with 2 µg/mL puromycin for three days. One day before use, cells were split into a 24-well (Cellvis; P24-1.5H-N) or 12-well (Cellvis; P12-1.5H-N) glass bottom plate. Cells were freshly transduced with lentivirus for each new experiment and used within 2 passages of transduction.

### Lentivirus construction

A custom vector for a codon-optimized HSV-1 ICP4 expression (based on KOS strain) was obtained from VectorBuilder (VB211009-1101fac, Addgene #250609). Codon-optimization was used to improve expression and better facilitate cloning. To construct the corresponding lentiviral vector (WT ICP4, pLH2488, Addgene #250611), the coding sequence was amplified and shuffled into a Tet-inducible lentivirus vector (pLVX-TetOn-puro, pLH2537, generously gifted by Meike Dittman) via Gibson assembly. To generate ICP4 mutant proteins(7, 23), primers were generated to amplify segments of ICP4 lacking either the C-terminal activation domain (ΔCTA, pLH2632, Addgene #250613) or the first 90 amino acids of the N-terminal activation domain (ΔNTA, pLH2631, Addgene #250612) and ligated between the EcoRI and BamHI sites of pLH2537. The DNA-binding domain mutant (R456L-R457L, mDBD)(28) was constructed from pLH2488 using overlapping primers containing the altered sequence, resulting in plasmid pLH2658 (Addgene #250614). All constructs were confirmed by DNA sequencing.

### Induction and chemical treatments

For all induction experiments, cells were seeded on a 24-well (Cellvis; P24-1.5H-N) or 12-well (Cellvis; P12-1.5H-N) glass bottom plate 24 h prior to use. For doxycycline (Dox) induction, lentivirus transduced or control cells were treated with 3 µg/mL dox for 9 h prior to imaging or collection for further processing.

### Immunoblotting

Total cellular protein was collected by lysis in sample buffer (62.5 mM Tris-HCl pH 6.8, 2% SDS, 10% glycerol, 0.7M β-mercaptoethanol) followed by boiling for 5 minutes. Lysates were fractionated by sodium dodecyl sulfate polyacrylamide gel electrophoresis (SDS-PAGE) and transferred to nitrocellulose membranes. Membranes were blocked in 5% non-fat milk in TBST for 1 h at room temperature and incubated in primary antibodies (ICP4: Abcam; ab6514, GAPDH: Cell Signaling; 2118S) at a 1:500 dilution in 3% BSA overnight at 4°C. Primary antibodies were detected using either anti-rabbit IgG HRP or anti-mouse IgG HRP secondary antibodies and visualized by chemiluminescent detection using an Invitrogen iBright FL1000 Imaging System.

### Immunofluorescence

Cells were treated as indicated, then immediately fixed at the indicated timepoints with 4% paraformaldehyde (Electron Microscopy Sciences, Cat. No. 15714) for 10 min at room temperature. The cells were subsequently washed three times with 1x PBS, and permeabilized with 0.5% Triton X-100 (Fisher Scientific, Cat. No. 9002-93-1) in 1x PBS for 15 min at room temperature. After blocking with 4% FBS in PBS for 1 h, primary antibody (ICP4: Abcam; ab6514, 1:500 dilution in 3% BSA) was applied overnight at 4°C. The next day, the cells were washed three times with 1x PBS and incubated with secondary antibodies (1:500 dilution) in 4% FBS in PBS at room temperature for 1 h in the dark. After washing three times with 1x PBS, cells were stained with SiR-DNA (Cytoskeleton; CY-SC007, 1:5000 dilution). Cells were subsequently stored in 1x PBS at 4°C in the dark until imaging. The fixed plates of cells were mounted on a Nikon spinning disk confocal scanning microscope. Fluorescence signals were obtained using 405 nm, 488 nm, 561 nm, and 640 nm lasers with the following emission filters: 455/50, 525/36, 605/52, and 705/72 (Chroma Technology Corp). Images were captured using a Prime 95B scMOS camera (Photometrics) with a 60x/1.49 numerical aperture objective lens with DIC capability.

### 5EU#incorporation assay

Cells were treated + / - Dox for 7 h, then exposed to a 200 uM 5EU pulse for 2 h to reach 9 h total of Dox treatment. Following EU exposure, cells were fixed with 4% PFA for 10 min at room temperature. The cells were subsequently washed three times with 1x PBS and permeabilized with 0.5% Triton X-100 in 1x PBS for 15 min at room temperature. Cells were then washed twice in 3% bovine serum albumin (BSA) (Fisher Bioreagents; BP1600-100) diluted in PBS. Cells were then biotinylated using the Click-iT Nascent Capture Kit (Fischer Scientific; C10365), following the manufacturer’s instructions. After completion of the Click-iT reaction, cells were blocked in 3% BSA for 1 h, after which primary antibody (ICP4: Abcam; ab6514, 1:500 dilution in 3% BSA) was applied overnight. Following overnight incubation with primary antibodies, immunofluorescence microscopy was performed as described above. To calculate 5EU mean intensity and background intensity, 20-μm Z-stacks were taken with 0.5-μm steps between frames, and images were analyzed with foci-counting(49).

### RNA isolation, cDNA synthesis, and RT-qPCR

RNA was extracted at the indicated timepoints using Ambion PureLink Mini Kit (Invitrogen; 12-183-018A) following the manufacturer’s protocol and after resuspension, resuspended RNA was stored at -80°C. WT and mutant ICP4 inductions were collected in biological triplicates, while mock-induced cells were collected in a technical triplicate.

### RNA-Seq library preparation and analysis

For RNA-Seq, extracted RNA samples were processed in the Genome Technology Center at NYU Langone Health. Sample quality was determined using a Bioanalyzer before automated stranded RNA-Seq library prep with polyA selection on ∼1 ug of RNA. Sequencing was performed on NovaSeq X+ 10B 100 Cycle Flowcells.

Paired-end sequencing reads (FASTQ format) of length 150 bp (ΔCTA/mDBD/ICP4 batch) or 50 bp (ΔNTA/ICP4 batch) were subjected to quality control using FastQC to assess read quality metrics(50). Adapter/quality trimming and filtering were conducted with Trim Galore (https://github.com/FelixKrueger/TrimGalore)(51) and minimum length and quality parameters adjusted after examining the FastQC report. Reads were aligned with STAR(52) to a modified GRCh38 reference genome that included the WT ICP4 or mutant ICP4 sequence. STAR was run with default parameters with output in sorted BAM format (--outSAMtype BAM SortedByCoordinate) and gene-level quantification enabled (--quantMode GeneCounts).

Mock controls were sequenced as a separate batch (paired-end length 50 reads). QC/Trimming/Alignment was performed as described above, aligning to unmodified GRCh38. Batch effect could not be modeled for our mock condition due to a lack of matching induced genotypes from the sequencing run.

Differential expression analysis was performed on all samples with a standard DESeq2 pipeline in R(53). Prior to DE analysis, lowly expressed genes were removed by filtering to those where at least 10 reads were mapped in a minimum of 2 or 3 samples, depending on group size. We report raw p-values, q-values and log2FC for all remaining genes. Linear models included a term to account for batch effect, except when not possible (siNTC comparisons). GSEA pre-ranked(35) was run for each DE comparison using the fgsea R package(54), ranking genes by sign(log2FC) x q-value.

For comparisons between our datasets and existing ICP4 Chip-Seq in WT HSV-1 infection, ChipSeq data including Input controls (FASTQ format) were downloaded from BioProject (<u>PRJNA553563</u>) and subjected to QC/Trimming/Alignment with FASTQC, Trim Galore and STAR (similar workflow as above), aligning to GRCh38 with KOS (HSV-1) genome appended. Peaks were called with MACS2(55) for each of 2, 4 and 6 h time points, producing narrowPeak files. The ChIPseeker R package was then used to annotate peak locations to genomic regions. Peaks were filtered to those found in Promotor and 5’ UTR regions and then overlap with DEGs from RNASeq analysis by the nearest gene.

For comparisons between our datasets and existing RNA-Seq in WT HSV-1 and n12 infection, Total RNA-Seq data including uninfected controls (FASTQ format) were downloaded from BioProject (<u>PRJNA851702</u>) and subjected to QC/Trimming/Alignment with FASTQC, Trim Galore and STAR (similar workflow as above), aligning to GRCh38 with KOS (HSV-1) genome appended.

Over-representation analysis using the R package clusterProfiler was performed for pathway enrichment tests involving unranked gene lists resulting from the intersection of data sets(38).

